# A 48-Channel Receive Array Coil for Mesoscopic Diffusion-Weighted MRI of Human *ex vivo* Brain Imaging on the 3T Connectome Scanner

**DOI:** 10.1101/2021.02.24.432713

**Authors:** Alina Scholz, Robin Etzel, Markus W May, Mirsad Mahmutovic, Qiyuan Tian, Gabriel Ramos-Llordén, Berkin Bilgiç, Thomas Witzel, Jason P Stockmann, Choukri Mekkaoui, Lawrence L Wald, Susie Y Huang, Anastasia Yendiki, Boris Keil

**Affiliations:** Institute of Medical Physics and Radiation Protection (IMPS), TH-Mittelhessen University of Applied Sciences (THM), Giessen, Germany; A.A. Martinos Center for Biomedical Imaging, Department of Radiology, Massachusetts General Hospital, Boston, Massachusetts, USA; Harvard Medical School, Boston, Massachusetts, USA; Harvard-MIT Division of Health Sciences and Technology, Cambridge, Massachusetts, USA; Center for Mind, Brain and Behavior (CMBB), Marburg, Germany

**Keywords:** Magnetic Resonance Imaging, Diffusion-Weighted Imaging, RF coil, receive array coil, brain imaging, *ex vivo* brain

## Abstract

*In vivo* diffusion-weighted magnetic resonance imaging is limited in signal-to-noise-ratio (SNR) and acquisition time, which constrains spatial resolution to the macroscale regime. *Ex vivo* imaging, which allows for arbitrarily long scan times, is critical for exploring human brain structure in the mesoscale regime without loss of SNR. Standard head array coils designed for patients are sub-optimal for imaging *ex vivo* whole brain specimens. The goal of this work was to design and construct a 48-channel *ex vivo* whole brain array coil for high-resolution and high *b*-value diffusion-weighted imaging on a 3T Connectome scanner. The coil was validated with bench measurements and characterized by imaging metrics on an agar brain phantom and an *ex vivo* human brain sample. The two-segment coil former was constructed for a close fit to a whole human brain, with small receive elements distributed over the entire brain. Imaging tests including SNR and G-factor maps were compared to a 64-channel head coil designed for *in vivo* use. There was a 2.9-fold increase in SNR in the peripheral cortex and a 1.3-fold gain in the center when compared to the 64-ch head coil. The 48-channel *ex vivo* whole brain coil also decreases noise amplification in highly parallel imaging, allowing acceleration factors of approximately one unit higher for a given noise amplification level. The acquired diffusion-weighted images in a whole *ex vivo* brain specimen demonstrate the applicability of the developed coil for high-resolution and high *b*-value diffusion-weighted *ex vivo* brain MRI studies.

## 1. Introduction

Diffusion MRI (dMRI) is a powerful, non-invasive technique for imaging axonal orientations as well as characterizing white and gray matter microstructure [1–5]. The basic premise of dMRI in the human brain is that the diffusion of water molecules in white matter is anisotropic, and that its preferential direction is aligned with the orientation of the underlying fibers [2]. A series of images, each sensitized to diffusion in a different direction, are acquired and used to infer the most likely orientation of water displacement in every voxel [6].

There are several requirements that increase the acquisition time needed for whole-brain dMRI. High spatial resolution is desirable for resolving small brain structures. A large number diffusion directions must be sampled to improve the angular resolution, *i*.*e*., the smallest angle between crossing fiber bundles that can be resolved. Finally, advanced dMRI sampling schemes may require images to be acquired with multiple *b*-values. Satisfying all these requirements would lead to acquisition times that are prohibitive for *in vivo* imaging in the absence of any image acceleration. As a result, trade-offs must be made that restrict *in vivo* whole-brain dMRI to the macroscale regime [4, 7], with voxel sizes on the order of 1 to 3 mm. Motion artifacts, which are exacerbated by long acquisitions, and distortions near tissue-air interfaces further degrade the effective resolution that is achievable *in vivo*.

Many of these issues can be circumvented in *ex vivo* dMRI, which allows for longer acquisition times, absence of motion and significantly reduced susceptibility artifacts with appropriate sample preparation [8]. Furthermore, *ex vivo* imaging enables the placement of coil elements closer to the actual brain tissue to maximize sensitivity. Thus, *ex vivo* imaging can achieve substantially higher spatial and angular resolution, permitting the anatomy and microstructure of complex fiber pathways to be imaged at the mesoscale, sub-millimeter regime, well beyond what is feasible *in vivo*. The impressive level of anatomical detail that can be resolved by *ex vivo* dMRI has already been demonstrated on a variety of human tissue samples [9–13]. *Ex vivo* dMRI, in combination with optical imaging, is an excellent tool for validating dMRI acquisition and analysis methods in human brain tissue [14, 15].

Higher spatial resolution comes at the cost of lower signal-to-noise-ratio (SNR). Several strategies for improving SNR in high-resolution *ex vivo* dMRI have been proposed and tested, mainly focusing on higher magnetic field strengths [16, 17], small-bore MRI scanners [18, 19] or high-performance gradient systems [20]. One of the main innovations introduced by the NIH Blueprint Human Connectome Project was the development of human scanners with ultra-high gradients, which allow high *b*-values to be achieved without loss of SNR [21]. Initial results have already shown the advantages of a 300 mT m^−1^ gradient system for imaging whole post-mortem human brains at 0.6 mm isotropic resolution [20], or smaller, non-human primate brain samples at 0.8 mm isotropic resolution [22]. Those results were obtained with an *in vivo* head coil. Dedicated *ex vivo* brain coils are known to increase signal reception sensitivity, and a few studies have shown the benefits of multi-channel brain array coils for *ex vivo* tissue imaging applications [23–25].

The aim of this study was to push the limits of spatial and angular resolution in *ex vivo* dMRI by designing, constructing, and validating a 48-channel (48ch) receive array coil for *ex vivo* whole human brain examinations. The array coil was developed for high spatial resolution and high *b*-value dMRI acquisitions with long scan times (a few hours to a few days) on the 3T Connectome scanner [20, 21]. This work presents high-sensitivity *ex vivo* diffusion MRI results obtained in a whole human brain specimen at mesoscale resolution (0.73 mm isotropic) using the 48ch receive coil on the 3T Connectome scanner and expands on preliminary results that were published in conference proceedings [26].

## 2. Material and Methods

### 2.1. Coil Design and Construction

To closely cover a whole human brain, we designed an anatomically-shaped *ex vivo* brain coil former (Figure 1 and 2) based on a nonlinear brain atlas of the International Consortium for Brain Mapping (ICBM). The coil housing was modeled with 3D computer aided design (CAD) software (Rhino3D, Robert McNeel & Associates, Seattle, WA, USA, version 6). It was designed to completely surround the brain with minimal space between the coil elements and imaging volume. The coil former was split into an upper and lower part, such that a whole brain can be placed inside the coil container. Both coil segments close with an overlapping rim structure. The coil container can accommodate whole brains with dimensions of 140 mm in the left-to-right direction, an anterior-to-posterior diameter of 182 mm, and a superior-inferior distance of 110 mm. The completed array coil is shown in Figure 3.

**Figure 1.**
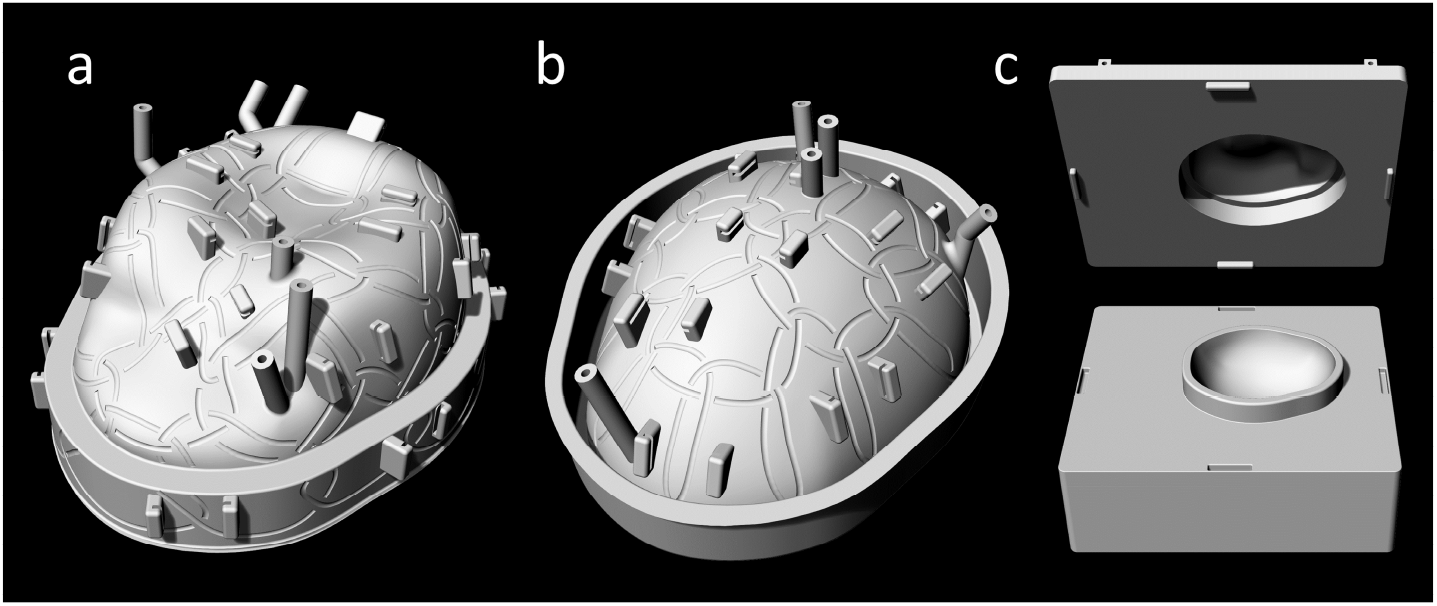
computer aided design model of the coilformer with graved loops and standoffs for preamplifier boards. **(a)** Top coilformer part **(b)** Bottom coilformer part **(c)** Inner side of both coilformer parts with overlapping frames to allow geometrical decoupling of the loops from top and bottom part.

**Figure 2.**
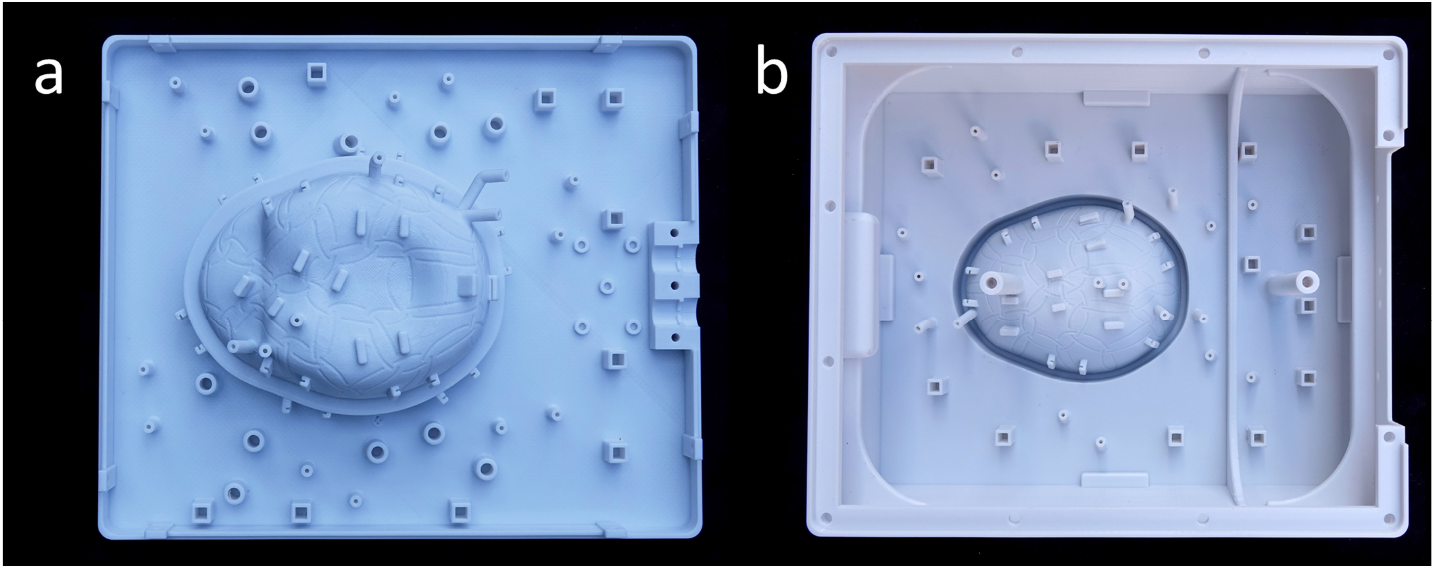
Polycarbonate printed coilformer with standoffs for preamplifier boards. **(a)** Top coilformer part. **(b)** Bottom coilformer part, in addition to two pillars to handle the weight of the brain and top part.

**Figure 3.**
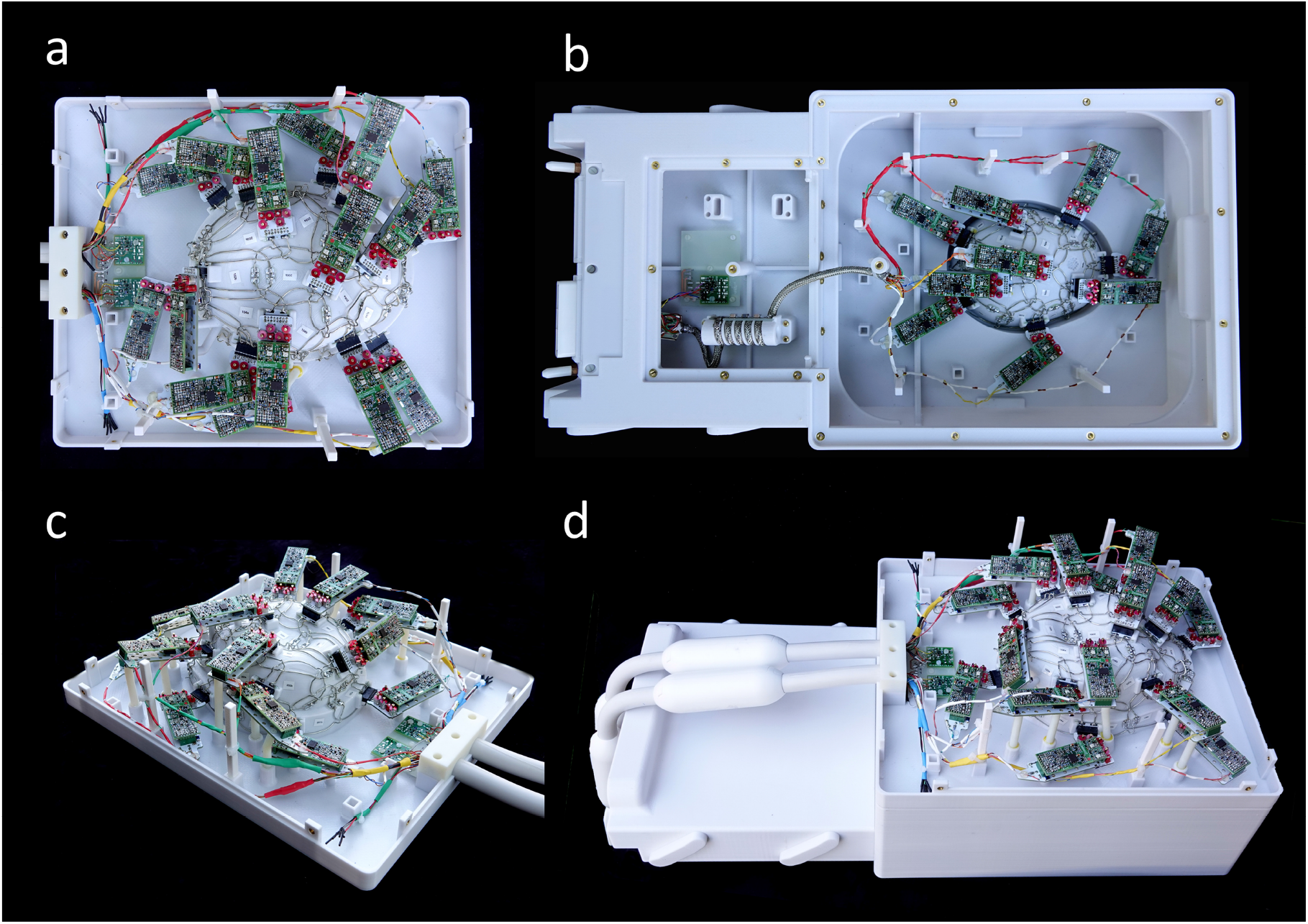
Completed coil with loops, preamplifier boards, preamplifiers, cable trap and EPROMs without coil covers **(a) and (c)** Top part with 30 coil elements **(b)** Bottom segment with 18 coil elements and a cable trap. **(d)** Assembled coil consisting of the top part and bottom part. The scanner connection cables from the top part involve two cable traps each and standard coil plugs. The bottom part uses a sliding connection mechanism.

In the bottom coil segment, we incorporated the mechanics for a plugging slide mechanism (Figure 3b and 3d), which directly plugs it into the scanner’s patient bed. The top coil segment is connected to the scanner using two standard multi-channel coil plugs. The *ex vivo* coil container was designed to allow the brain to be placed at the isocenter of the scanner.

The positions of the 48 coil elements on the outer surface of the coil former were derived from a hexagonal/pentagonal tiling pattern [27], with 30 and 18 coil elements located on the top and bottom segments, respectively (Figure 4).The position and outline of all loop elements, which are decoupled geometrically from neighboring loops by critical overlap [28], were incorporated in the CAD model. The majority of the loops was circularly formed, whereas some loops were arbitrarily shaped to fit over the rim structure. The loop size and thus the critical overlap was determined empirically in previously tested bench measurements. The average diameter of the circular coil elements is 54 mm with an inductance of about 203 nH. The overlap of these loops is about 0.27 times the loop diameter. Standoffs for circuit boards and cable routing were implemented to provide stable mounting positions. The coil former including its cover were then 3D-printed in polycarbonate (PC) using a 3D printer (Fortus 350, Stratasys, Eden Prairie, USA).

**Figure 4.**
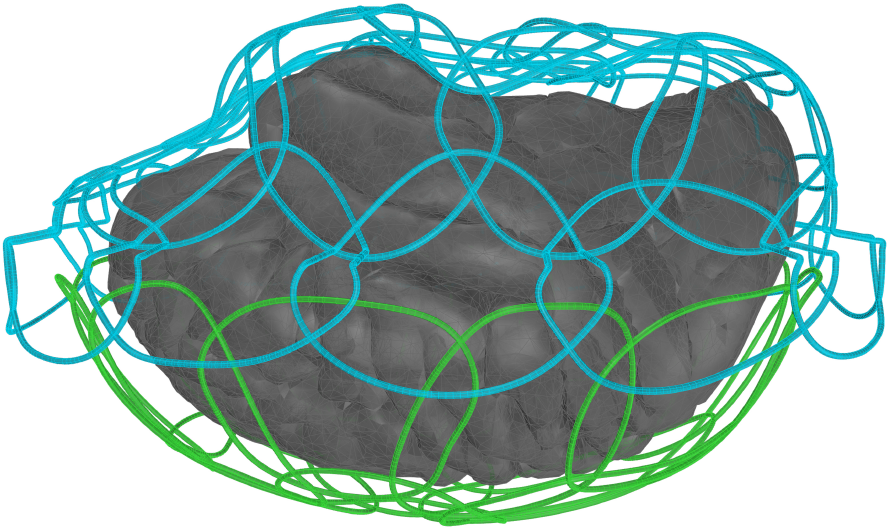
Placement of the 30 top loops (blue) and the 18 bottom loops (green) around the brain (gray) in the computer aided design program.

### 2.2. Coil Circuit

The loop elements were constructed out of 1.3 mm thick tin-coated copper wire. Compared to flat circuit board copper traces, the wire loops reduce eddy current losses in a high-density array coil architecture [29]. Implemented small bridges in the conductor enable one loop to cross over another without touching [30].

Each coil circuit (Figure 5) consists of a loop with three symmetrically placed ceramic capacitors (Series 11, Voltronics, Danville, NJ), one variable plastic capacitor (GFX2700NM; Sprague Goodman, Westbury, New York, USA), a matching network to the preamplifier (Siemens AG, Healthineers, Erlangen, Germany), and an actively controllable detuning resonant circuit. A typically redundant passive detuning safety mechanism for *in vivo* examinations was omitted for this *ex vivo* coil.

**Figure 5.**
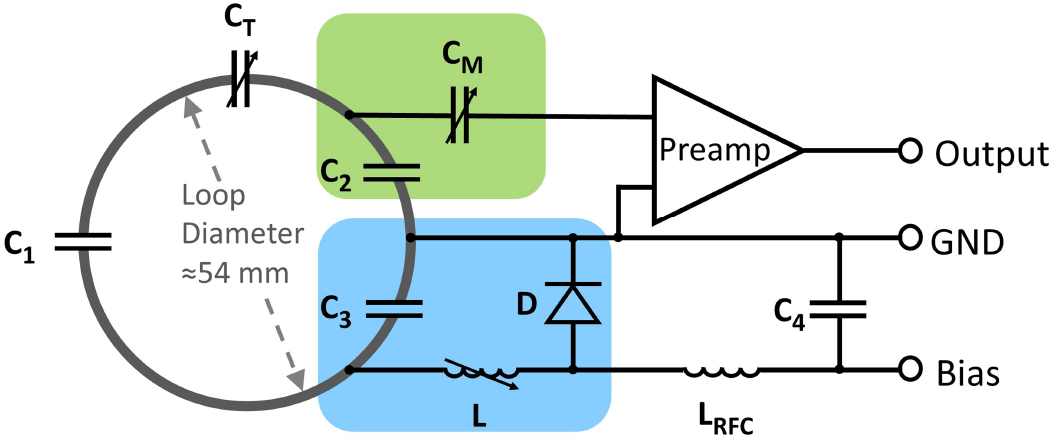
Circuit schematic for one coil element. Each loop consists of three fixed capacitors (*C*_1_-*C*_3_) and one variable capacitor (*C*_*T*_). *C*_*T*_ fine-tunes the resonant frequency of the coil to Larmor frequency corresponding at 3T. *C*_2_ and *C*_3_ create a capacitive voltage divider. *C*_3_ is part of the active detuning circuit (blue) together with the variable inductor *L* and the PIN diode *D. C*_2_ and *C*_*M*_ (green) provide both impedance matching of the loop and impedance transformation to establish preamplifier decoupling. Typical values for the components are: *C*_1_ = 33 pF, *C*_2_ = 56 pF, *C*_3_ = 56 pF, *C*_*T*_ *≈* 18 pF, *C*_*M*_ *≈* 18 pF, *L ≈* 24.5 nH, *L*_*RF C*_ = 2.7 µH.

The variable capacitor *C*_*T*_ (3 −33 pF) was used to fine-tune the loop resonance to the Larmor frequency at 3T (123.25 MHz). *C*_2_ and *C*_3_ create a capacitive voltage divider. The variable capacitor *C*_*M*_ (3 −33 pF, GFX2700NM; Sprague Goodman, Westbury, New York, USA) provides impedance matching of the loop output to a 50 Ω noise matched condition needed by the preamplifier to operate at the lowest noise figure at 123.25 MHz) [31]. To ensure accurate detuning of the loop elements, an active detuning circuit was implemented. It consists of one of the voltage dividing capacitors *C*_3_ and a variable inductor *L* (Coilcraft Inc., 25 −32 nH, 165-02A06L, Cary, IL, USA) in series to a PIN diode *D* (MA4P4002B-402; Macom, Lowell, MA, USA) [32]. During transmit, a DC current is applied to forward bias the PIN diode. This in turn activates the detuning resonant circuit at the Larmor frequency and generates a high impedance in the loop to suppress current flow. The RF-choke *L*_*RF C*_ (Coilcraft Inc, 2.7 µH, 1812CS-333XJLC Cary, IL, USA) and *C*_4_ block the RF signal to prevent passing into bias source.

While nearest neighbors use geometrical decoupling, next-nearest neighbors and further coil elements are decoupled by the impedance transformation of the input of the preamplifiers [28]. The capacitors *C*_2_ and *C*_*M*_ and the preamplifier’s input impedance form a resonant circuit, which enables a voltage-source measurement setup, where RF current flow is minimized. As a consequence, inductive coupling across elements is highly reduced and all coil elements receive independently, while maintaining a 50 Ω output impedance.

Both the matching and detuning network of the coil element are placed on the preamplifier’s daughter board, rather than soldering these components directly to the coil former. Therefore, the daughter board is a part of the coil element. The printed circuit board (PCB) daughter board is connected to the loop with an intermittent pin connector. This setup allows a fast construction process of dense array coils.

According to the RF scanner architecture, pre-amplified signals from two loops elements are multiplexed onto one output coaxial cable. The bundled output cables are passed through cable traps to prevent RF common mode currents on the shield of the coaxial cable [33]. The cable traps comprise a wounded coaxial cable bundle, which form an inductance (*≈* 109 nH), and a parallel ceramic high power capacitor (15.2 pF, Series 25, Voltronics, Danville, NJ), which resonates at Larmor frequency. Two traps are incorporated into the cables of the upper array coil segment and one cable trap is located directly in the bottom coil housing part.

### 2.3. Coil Bench Measurements

For bench measurements during the construction process, a custom-made coil plug simulator was used. It provides voltage for the preamplifiers (3 V) and the opportunity to apply a DC current (100 mA) to bias manually each PIN diode forward, which allows for active detuning of single coil elements. To gather information about bench level metrics, e.g. transmission and reflection measurements, a vector network analyzer (VNA) (ENA series, Agilent Technologies, Santa Clara, CA) and custom-built RF tools such as single / double probes and sniffer probes were used. These measurements included tuning to Larmor frequency, active detuning, preamplifier decoupling and geometrical nearest neighbor decoupling of each coil element.

The loops were tuned under a *S*_21_ control with a 50 Ω dummy load plugged into the preamplifier socket, while all other elements of the array were detuned. Active detuning was performed by using *S*_21_ measurement with the double-probe for each loop, while all other coil elements were detuned and the relevant loop under test was switched between the tuned and detuned state. The difference of both states at the Larmor frequency indicates the magnitude of active detuning. A similar *S*_21_ double-probe measurement was carried out to determine the effectiveness of the implemented preamplifier decoupling, first by plugging the preamplifier into the socket on the PCB and second by terminating the socket with a load impedance of 50 Ω. Again, all but the loop element to be tested were detuned.

Coupling of nearest neighbor elements was measured with direct *S*_21_ VNA measurement by using coaxial cables, which were directly plugged into the preamplifier sockets. During this measurement, all other coil elements were detuned. This measurement configuration was also used to verify 50 Ω coil impedance matching using *S*_11_ and *S*_22_ measurements. [30, 31]

Furthermore, unloaded-to-loaded coil quality factor ratio (*Q*_*U*_ */Q*_*L*_) of one representative coil element was measured within the populated but detuned array assembly, using the *S*_21_ double-probe method [34]. As a load, a fixed tissue brain sample in periodate-lysine-paraformaldehyde (PLP) solution was used.

### 2.4. MRI data acquisition and analysis

Imaging metrics were acquired on a clinical 3T MRI scanner (MAGNETOM, Skyra, Tim 4G, Dual Density Signal Transfer, Siemens AG, Healthineers, Erlangen, Germany), equipped with a customized gradient coil (AS302 CONNECTOM 1.0 gradient)^1^ with a maximum gradient strength of 300 mT m^−1^ and a maximum slew rate of 200 T m^−1^ s^−1^.

For evaluating the developed *ex vivo* whole brain array coil, we constructed a human-brain-shaped phantom using a 3D printer (Objet30 Pro, Stratasys, Eden Prairie, USA). The phantom was filled with a mixture of 830 ml distilled H_2_O, 29 g NaCl, 12.5 g of agar powder (Sigma-Aldrich Corp., St. Louis, MO) and 936 g sugar [35]. The electrical characteristics of the brain phantom were determined with a VNA equipped with a dielectric probe kit (85070E kit, Agilent Technologies, Santa Clara, CA) and were representative of the averaged human brain (*e*_*r*_ = 66.34 and *σ* = 0.49 S m^−1^). For determining SNR and G-factor, the phantom was scanned with a proton density (PD)-weighted FLASH sequence (repetition time (TR) = 200 ms, echo time (TE)= 4.8 ms, flip angle (F) = 15°, matrix (M): 192 × 192 (SNR) and 64 × 64 (G-factor and SNR in parallel imaging), field of view (FOV): 256 ×256 mm^2^, slice thickness: 8 mm, bandwidth (BW): 200 *Hz/pixel*). Information about noise correlation was obtained with the same sequence but without RF excitation.

Pixel-wise SNR maps were calculated using the noise-covariance-weighted, root sum-of-squares image reconstruction method from Kellman *et al*. [36]. To evaluate the array coil’s encoding capability for parallel imaging, SENSE G-factor maps were computed using the acquired noise correlation matrix and complex sensitivities of the coil elements [37]. The FOV of the G-maps was tightly enclosed to the phantom, in order to enhance the aliasing pattern inside the imaging object.

A valuable metric is the remaining image SNR after the parallel imaging acceleration has been performed. We calculated the remaining SNR by dividing the SNR globally by the square root of the reduction factor *R* and further locally with the noise amplification given by the G-factor.

For further characterization of the coil performance, we examined the encoding power for simultaneous multislice (SMS) acquisitions with blipped-controlled aliasing in parallel imaging [38–40]. To assess the encoding capability of combined SMS and in-plane acceleration, a reduction factor of *R* = 2 and a slice acceleration factor from *MB* = 4 up to *MB* = 8 with a 1/3 FOV shift were evaluated. Noise correlation and SNR and G-factor maps of the 48ch *ex vivo* brain coil were compared to a customized 64-channel (64ch) whole head receive array coil [41] with identical acquisition parameters.

In addition, time course stability of each coil element was measured with a single-shot gradient echo echo planar imaging (EPI) sequence (time points: 500, TR = 1000 ms, TE = 30 ms, F = 90°, M: 64 × 64, FOV: 200 ×200 mm^2^, slices: 16 slices of 15 mm, BW: 2298 Hz px^−1^) with the brain phantom. The average intensity of a 15-pixel square region of interest (ROI) in the brain center was de-trended with linear and quadratic temporal trends and plotted. The stability was calculated as the variation of signal intensity from peak-to-peak as a percentage from the average signal intensity[42].

Finally high-resolution (0.73 mm isotropic) diffusion imaging was performed on a whole, *ex vivo* human brain. The brain had been excised from a male who had died of non-neurological causes, and had been placed in fixative (10% formaldehyde) for 90 days before being transferred to paraformaldehyde-lysine-periodate solution for long-term storage. Diffusion-weighted images were acquired with a 3D diffusion-weighted spin echo echo planar imaging multi-shot sequence (TR = 500 ms, TE = 65 ms, echo spacing: 1.22 ms, M: 160 × 268 × 208, FOV: 117 ×196 mm^2^, BW: 1244 Hz px^−1^, 16 shots, EPI factor= 10, no partial Fourier, maximum gradient strength of 91 mT m^−1^). One image with *b* = 0 and 16 diffusion-weighted volumes with *b* = 4000 s mm^−2^ and non-colinear diffusion encoding directions were acquired. Diffusion-weighted volumes were corrected for eddy current distortions with the *eddy* tool from FSL [43]. The diffusion tensor model was fit by linear least-squares fitting of the logarithm of the dMRI signal with the *dtifit* tool in FSL. The phase-encoding direction was anterior-posterior when considering the conventional sagittal plane. Since the brain in the constructed coil was rotated compared to the usual orientation of a patient, the anatomical axis of the phase-encoding direction was inferior-superior.

## 3. Results

### 3.1. Coil Bench Measurements

The *Q*_*U*_ */Q*_*L*_-ratio of a 54 mm loop element was measured to be 233*/*46 = 5.1 with six surrounding but non-resonant neighboring loops. Thus, the array’s loop elements operate in the sample noise dominated regime. The geometrical decoupling of nearest neighbors was *S*_21_ measured with an average value of −16 dB and ranged from −14 dB to −18 dB. Non-adjacent and thus non-overlapping coil elements, which are primarily decoupled via preamplifier decoupling, obtained an average decoupling value of −18 dB with a range from −17 dB to −19 dB. The isolation between tuned and detuned states caused by the active detuning circuit reached an average value of 42 dB.

### 3.2. Image Performance

Figure 6 shows the noise correlation matrix of the 48ch *ex vivo* coil and that of the 64ch *in vivo* coil. The *ex vivo* array has a range of noise correlations from 0.02 % to 35.8 % with an average value of 7.5 %, while the *in vivo* array has noise correlations from 0.12 % to 53.8 % with an average value of 7.1 % for the off-diagonal elements.

**Figure 6.**
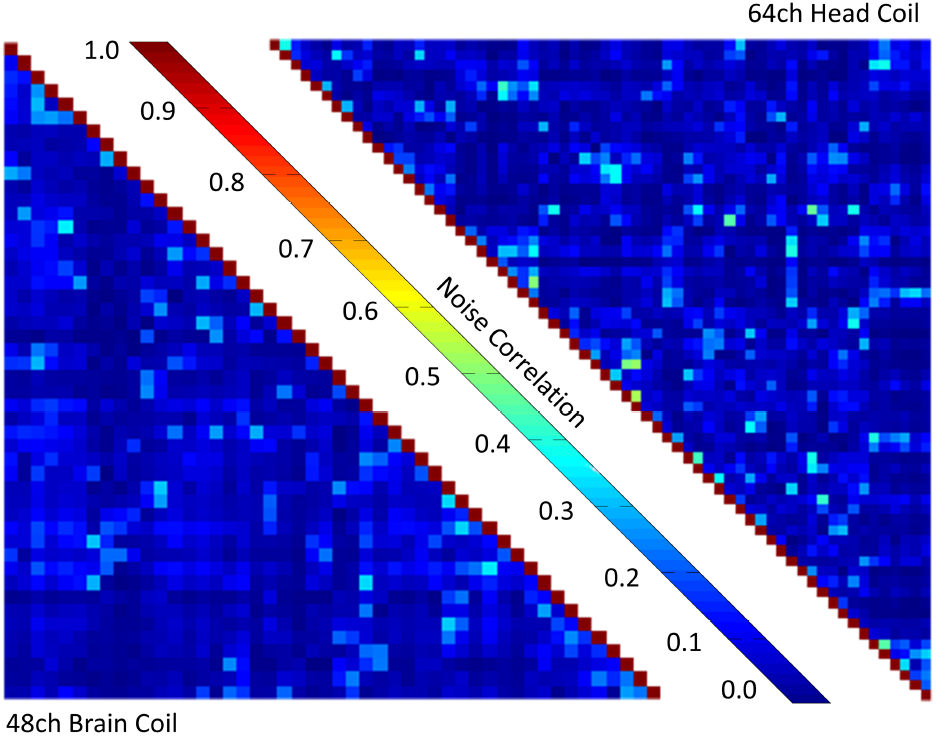
Noise correlation matrix of the 48ch *ex vivo* brain coil and the 64ch *in vivo* head coil with the scale normalized to 1.

Figure 7 compares the SNR maps from the newly developed 48ch *ex vivo* brain coil to that of the existing, custom 64ch whole head coil, in different planes of the agar phantom. For both coils, the measured SNR is highest in the outer periphery and decreases towards the phantom center. The newly constructed 48ch array coil outperforms the larger 64ch *in vivo* head coil by a factor of 2.5, when the average SNR over the whole brain phantom volume is considered. In the periphery and in the center of the phantom, a 2.9-fold and 1.3-fold SNR gain was measured, respectively.

**Figure 7.**
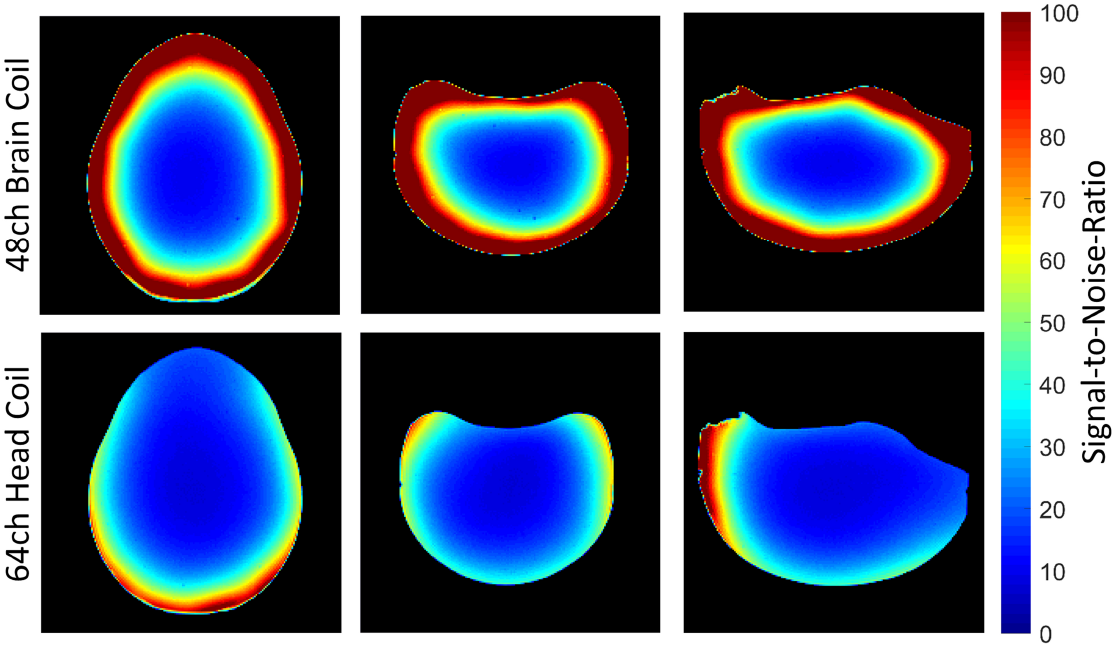
Comparison of the signal-to-noise-ratio, normalized to 100, of a transverse (left), coronal (middle) and sagittal (right) slice of the brain phantom with the 48ch *ex vivo* brain coil (top row) and the 64ch head coil (bottom row). The 48ch *ex vivo* brain coil shows a 1.3-fold SNR gain in the center and a 2.9-fold SNR improvement in the peripheral regions when compared to the 64ch head coil.

Figure 8 shows the SENSE inverse G-factor maps in a representative coronal plane of the brain phantom for both one-dimensional and two-dimensional acceleration obtained from the 48ch *ex vivo* brain coil and the 64ch *in vivo* head coil. The newly constructed 48ch coil provides significant improvement compared to the 64ch head coil for both in-plane acceleration types. Both coils show minimal noise amplifications for acceleration factors of *R* = 2, *R* = 3 and *R* = 2×2. However, for higher accelerations (*R >* 3) the 48ch *ex vivo* coil provides favorable encoding capabilities when compared to the 64ch *in vivo* head coil. At *R* = 4, the 48ch coil shows on average a 16 % lower G-factor than the 64ch head coil. When comparing the peak G-factors between both, the 48ch coil shows a 21 % improvement. The enhanced encoding power of the 48ch coil becomes even more apparent when very high acceleration factors are compared. The improved average and peak Gfactor for *R* = 7 was measured to be 35 % and 41 % lower. At *R* = 5×5 the noise amplifications could be reduced on average by 43 %, while the peak G-factor decreased by 53 %.

**Figure 8.**
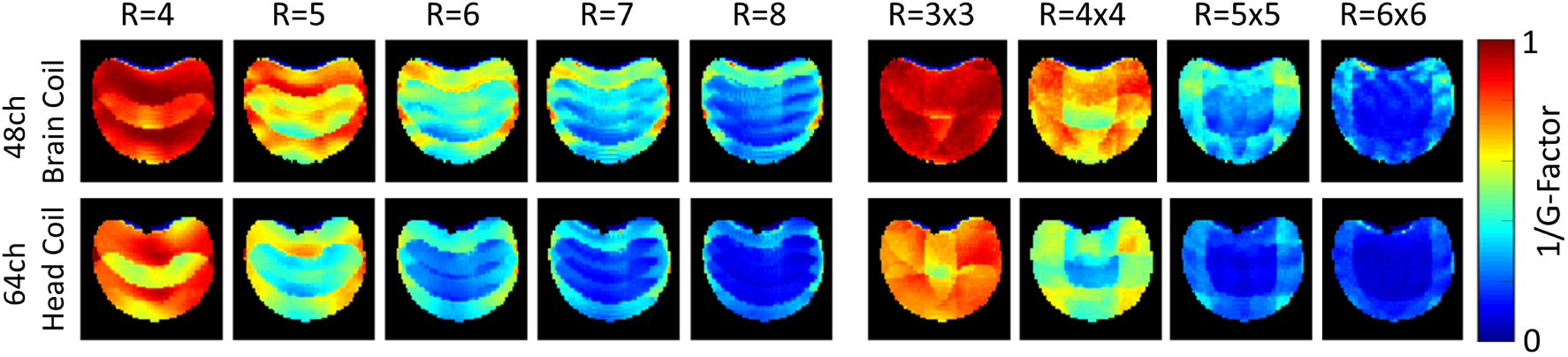
Comparison of inverse phantom G-factor maps between the 48ch *ex vivo* brain coil (top row) and the 64ch head coil (bottom row) for different acceleration factors (*R*) obtained from representative coronal slice. The G-Factors from the 48ch *ex vivo* brain coil show overall lower noise amplification, when compared to the 64ch head coil.

A more meaningful figure of merit is the SNR obtained from the accelerated image, where both the under-sampled *k*-space trajectory and the local noise amplification were taken into account. Figure 9 illustrates the accelerated SNR for both coils using box plots and its corresponding average values. Since the constructed 48ch coil provides both a higher baseline SNR and lower G-factors, it highly outperforms the 64ch head coil across all acceleration scenarios. The average SNR from the 64ch coil only reaches the lower 25th percentile of the 48ch ex vivo coil. Further, it should be noted that the relative gain in average SNR increases with higher acceleration factors (*e*.*g*., factor 2.4 for *R* = 2 and 3.9 for *R* = 7) for both one-dimensional and two-dimensional acceleration.

**Figure 9.**
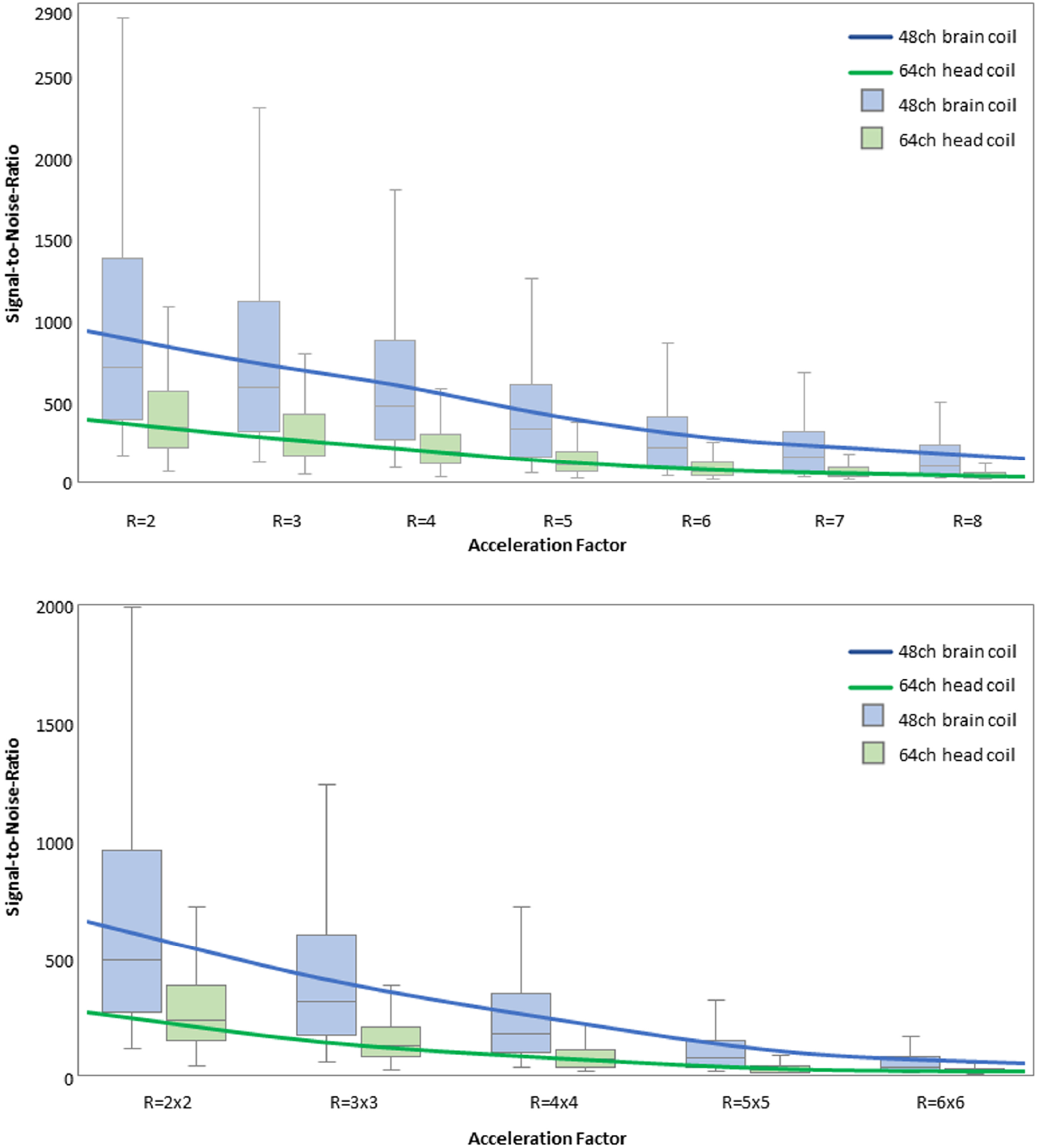
Parallel imaging accelerated signal-to-noise-ratio as a function of acceleration factor (*R*) from the 48ch brain coil and the 64ch head coil for one-dimensional (top) and two-dimensional (bottom) accelerations. The continuous lines indicate the average SNR, box plots represent median (horizontal line), lower/upper quartiles and minimum-maximum range (whiskers) without outliners. The constructed 48ch coil shows higher accelerated SNR in the entire range of acceleration factors.

Figure 10 compares the inverse G-factor maps for the SMS image reconstruction technique from a coronal slice of the brain phantom. Compared to the 64ch head coil, the constructed 48ch coil indicates overall substantially lower noise amplification for the SMS examination, as well as for combined SMS and in-plane acceleration. At a multiband factor of *MB* = 4, the 48ch coil generates negligible noise amplifications (*g*_mean_ = 1.0002 and *g*_max_ = 1.0569), while the 64ch head coil shows substantial noise gains of *g*_mean_ = 1.1218 and *g*_max_ = 1.6345. Furthermore, the dedicated 48ch *ex vivo* brain coil achieves similar to slightly better encoding capabilities at *MB* = 8 as the 64ch head coil at *MB* = 4 (*g*_mean48ch_ = 1.0047 *vs. g*_mean64ch_ = 1.1317 and *g*_max48ch_ = 1.2913 *vs. g*_max64ch_ = 1.6345). Therefore, the 48ch coil allows the application of a slice acceleration factor of *MB* = 8 with negligible noise gain.

**Figure 10.**
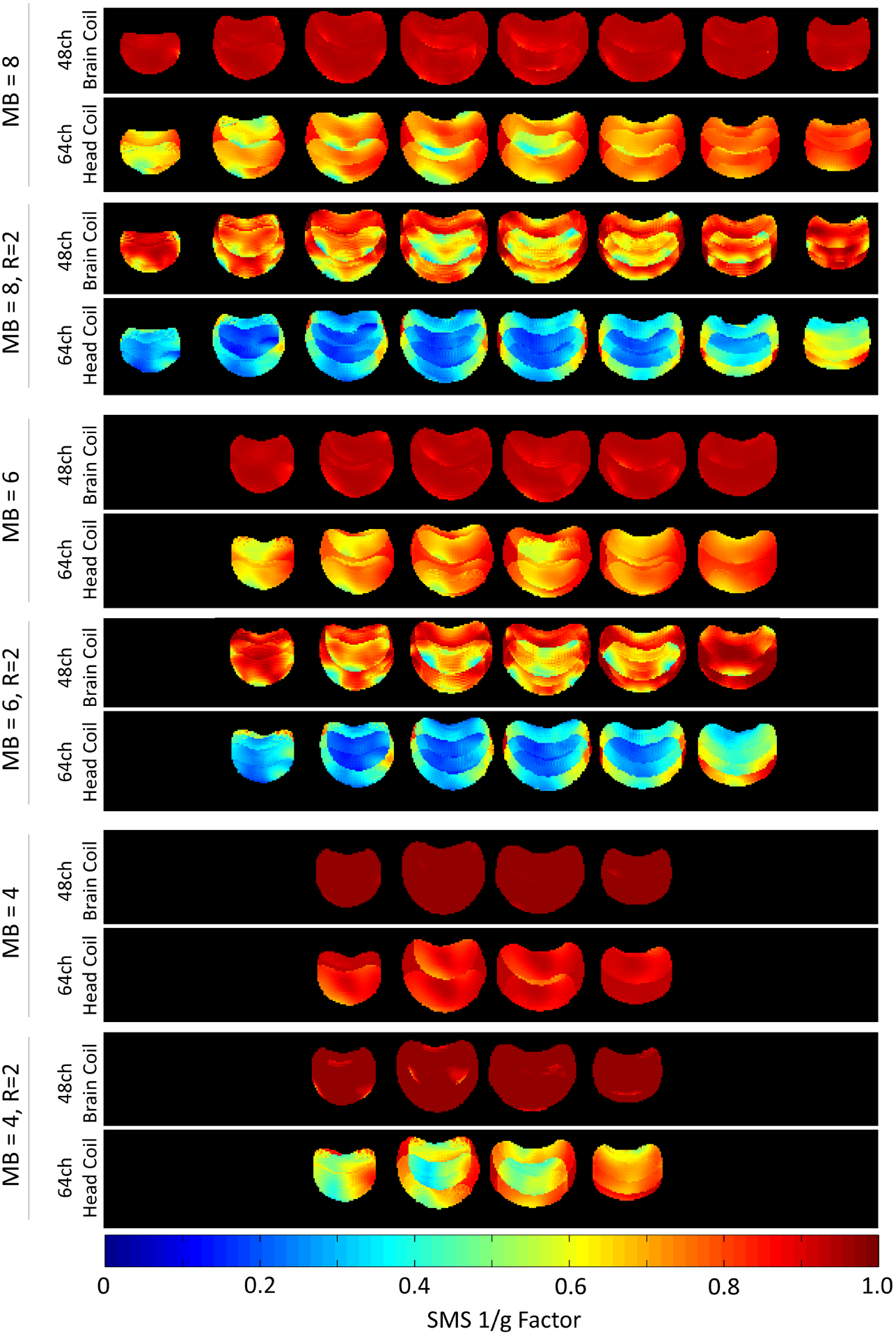
Comparison of inverse G-factor maps of the brain phantom for accelerated imaging with simultaneous multislice technique. The 48ch brain coil shows overall considerable lower noise amplification in comparison the the 64ch head coil.

To assess the accelerated SNR during SMS acquisitions, Fourier averaging needs to be taken into account: In the case of the *MB* = 8 acceleration, eight times more ^1^H spins are simultaneously excited compared with a single-slice acquisition. Thus, for a multiband factor *MB*, the SNR efficiency can be improved up to a factor of √*MB*. This translates to an SNR increase by a factor of up to 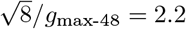, when compared to a commonly used consecutive single-slice acquisition schemes. The *MB* = 8 achievable SNR obtained from the 64ch is only increased by a factor of up to 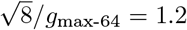. In direct comparison, when the baseline SNR, Fourier averaging, and G-factors are taken into account, the 48ch coil achieves up to a 4.5-fold SNR improvement at *MB* = 8 compared to the 64ch head coil. Time course stability tests show a peak-to-peak variation of 0.4 % over 500 time-points EPI sequence measured in a ROI comprising 15 × 15 pixels.

As an initial demonstration of the high sensitivity images that can be acquired in *ex vivo* whole brain specimens using the 48ch coil, maps obtained from a 0.73 mm isotropic resolution dMRI scan of a whole fixed brain are shown in Figure 11. The five columns show: the *b* = 0 image, a diffusion-weighted image (left-right diffusion-encoding direction), the mean diffusivity map, the fractional anisotropy (FA) map, and the FA map color encoded by the principal eigenvector (V1) of the diffusion tensor. The figure includes a coronal view (a), with a magnified area showing fine anatomical detail in the striatum (b), and an axial view (c), with a magnified area showing radial fibers in the primary motor cortex (d). These maps illustrate the capability of our *ex vivo* coil to map detailed circuitry both in deep brain and near the cortical surface and demonstrate the feasibility of obtaining data at high spatial resolution using the high gradient strengths available on the 3T Connectome scanner.

**Figure 11.**
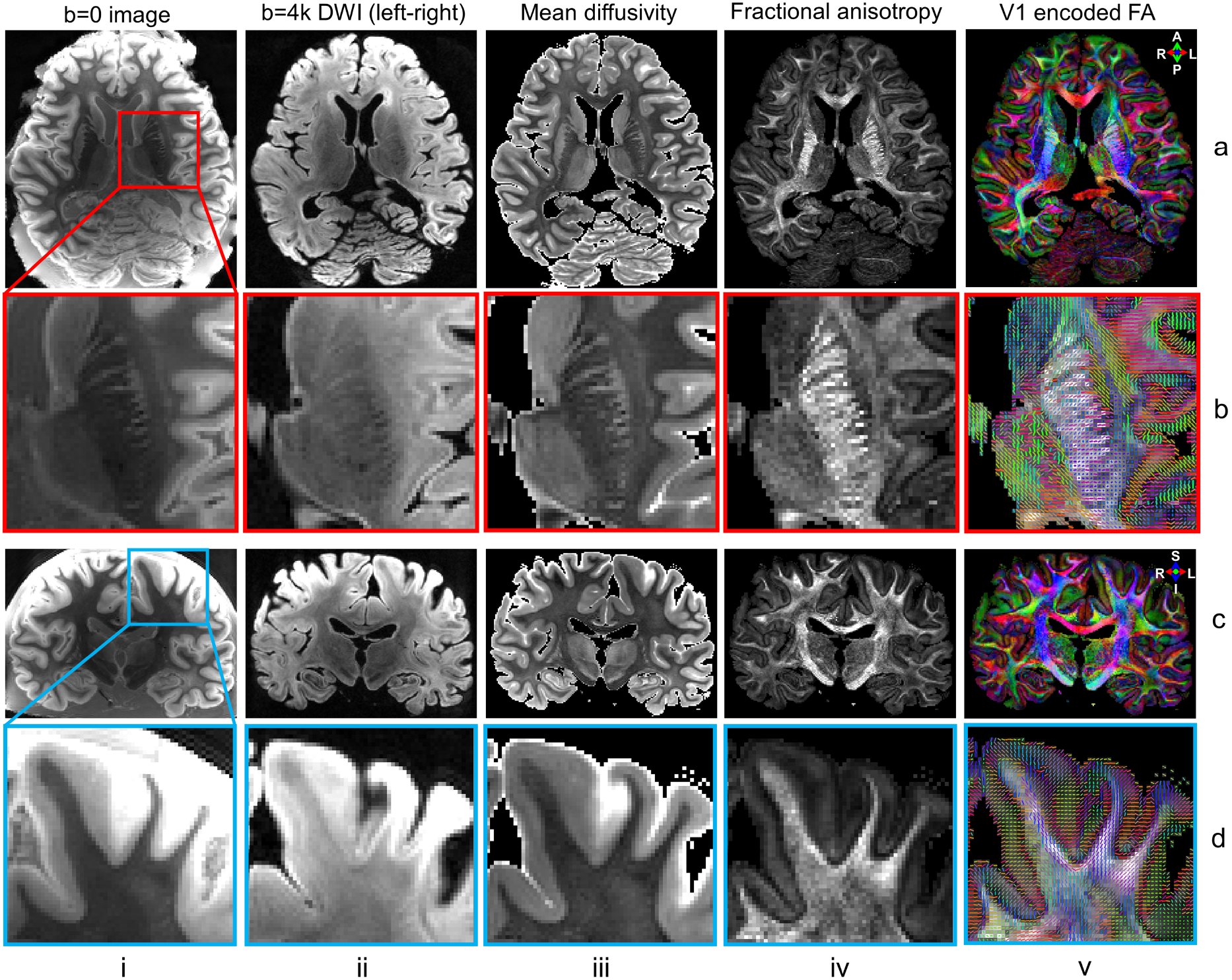
High-resolution Diffusion Tensor Imaging (DTI) results at 0.73 mm isotropic resolution with *b* = 0 images (column i), diffusion-weighted images (DWI) acquired at *b* = 4000 s mm^−2^ along left-right diffusion-encoding direction (column ii), mean diffusivity maps (column iii), fractional anisotropy (FA) maps (column iv) and FA maps color encoded by the primary eigenvectors (V1) from DTI (column v). Two regions of interest in the deep white matter (red box) and sub-cortical white matter (blue box) are displayed in enlarged views (rows b and d) showing fine-scale structures in the internal capsule and cerebral cortex.

## 4. Discussion

We designed, constructed, and evaluated a 48ch *ex vivo* brain array receive coil for high-resolution and high *b*-value dMRI of a whole *ex vivo* human brain on the 3T Connectome scanner [20, 21]. The coil was characterized by both bench tests and image metrics. Bench tests included element measurements of the coil quality factor *Q*, active detuning, geometrical decoupling, and preamplifier decoupling. MRI evaluations included measurements of the noise correlation, pixel-wise SNR, and G-factor, as well as time course stability using a brain shaped agar phantom. We demonstrated the coil’s performance in achieving high SNR with the acquisition of 0.73 mm isotropic resolution diffusion-weighted MR images of a whole *ex vivo* brain.

In many applications, large channel count arrays with relatively small loop sizes such as the 54 mm loops used here are necessary to increase both reception sensitivity and encoding power. However, very small loop elements quickly lose their sample noise dominance. Under these circumstances, small elements do not translate to higher SNR acquisitions anymore. For *in vivo* imaging at 3T, this critical size is reached at about 60 mm diameter [44]. In *ex vivo* brain imaging, however, loop sizes can be made substantially smaller than for *in vivo* imaging. This is attributed to the brain fixation medium, which has a higher conductivity compared to *in vivo* tissue and thus provides a higher fraction of sample noise. While the noise increases in the *ex vivo* sample, the electronic noise can be decreased by omitting *in vivo* human safety features in the coil element circuity, such as passive detuning and RF-fuses. This condition results in an enhanced *Q*_*U*_ */Q*_*L*_-ratio when using small receiver elements. Therefore, the implemented loop size of *≈* 54 mm provides a relatively high *Q*_*U*_ */Q*_*L*_-ratio of 5.1, outperforming most coils optimized for *in vivo* applications with loop diameters ranging from 50 mm to 65 mm from our previous studies [41, 44, 45]. As a consequence, the minimum loop diameter at which sample noise dominance is maintained decreases for imaging fixed tissue brain samples in PLP solution, allowing us to contemplate very high-density arrays for *ex vivo* sample examinations.

Despite RF electrical optimizations, the mechanical coil former is an important and critical design aspect for *ex vivo* imaging. To improve SNR, the loops were populated very close to the sample, maximizing signal reception. Thus, the completely brain-enclosing coil former with uniformly distributed loop elements guarantees nearly omni-directional signal reception from the sample (decreased sensitivity was observed with some loops aligned to be almost parallel to the magnetic field *B*_0_). However, an entirely surrounding coil array requires a split housing mechanism, which disturbs the loop layout and makes it difficult to maintain geometric decoupling at the split housing edge. Therefore, an overlapping edge structure was implemented, enabling adjacent loop elements to be geometrically decoupled across the two housing segments, while the overall array coil structure remains self-contained.

In array coil design, the central ultimate SNR is already approached with only 12 surrounding coil elements at 3T [46]. Implementing higher loop element counts only yields SNR improvements at the periphery for a given geometry. Nevertheless, relative central SNR gains are achievable with tightly fitting array coils. Due to the lack of dedicated *ex vivo* receiver arrays, *in vivo* head coils are commonly used in many *ex vivo* brain studies [3, 47–49]. However, these coils are not well suited in terms of sample fitting and SNR performance. Optimizing both the mechanical features for close fitting of samples and the RF circuitry can thus result in significant SNR gains in the brain. This implementation provides a 30% SNR increase of the 48ch coil at the phantom center when compared to the larger 64ch head coil. In addition, in the peripheral regions of the brain phantom, the tight-fitting form factor also provides favorable SNR gains, as evidenced by an almost 3-fold SNR improvement over the 64ch coil. The high SNR can be exploited to reduce the voxel size, enabling high spatial resolution MR imaging of a whole *ex vivo* brain.

The average noise correlation of 9 % indicates a well decoupled array and highly independent operating receiver loops. Adjacent loops show much higher coupling values up to 36 %, which can be attributed to insufficient overlap, resulting in a remaining mutual inductance and shared resistance especially in the sample voxels beneath the overlapping loop regions.

The constructed 48ch *ex vivo* brain coil shows remarkably better encoding performance when compared to the 64ch head coil. The encoding power of the 48ch coil enables approximately one additional acceleration unit, for both one-dimensional and two-dimensional accelerations, with the same noise amplification as the 64ch head coil. Improvements in G-factors are usually achieved by implementing higher channel counts on a given geometry. However, when comparing array coil formers of different sizes, similar improvements in G-factors can be achieved by (1) reducing the diameter of the coil elements at constant or even lower channel counts, and (2) positioning the coil elements in close proximity to the sample. The tight-fitting, smaller loop elements of the constructed 48ch coil provide an overall stronger spatial modulation in the signal sensitivity’s magnitude and phase. Consequently, this coil arrangement allows favorable encoding capabilities for unaliasing folded images (SENSE method) or synthesizing spatial harmonics (GRAPPA or SMASH methods). Additionally, the entirely enclosed *ex vivo* coil former of the 48ch coil leads to better spatial coverage for the aliased pixels when compared to a head array coil, which obviously has limited coverage along the inferior aspect and in the area covering the face.

Reducing scan time using parallel imaging techniques is not strictly essential when constraints on acquisition time are lifted for *ex vivo* examinations. On the other hand, 2D acquisitions are still often used despite their SNR inefficiency per unit time [11, 22]. For example, mapping tissue microstructural features such as axon diameter throughout the whole human brain involves measurements at multiple *b*-values [50], and protocol optimization may be facilitated by 2D scans acquired at resolutions on the order of 0.8 to 1 mm isotropic. For such 2D acquisitions, slice acceleration enables the excitation and measurement of multiple slices [39, 40, 51]. Unlike conventional parallel imaging, which requires under-sampled data acquisition, these techniques provide acceleration by exciting the spins in multiple slices at the same time using multi-band radiofrequency pulses. These newer multi-band MR acquisitions have the SNR advantages of 3D sampling based on Fourier averaging [38, 39]. Therefore, SNR efficiency can be improved by up to a factor of √*MB*. In practice, however, the SNR gain is reduced locally by the SMS G-factor of the coil and by changes in the sequence parameters, *e*.*g*., reduction of TR. The SNR recovery achieved by the SMS method is highly advantageous for dMRI, which normally suffers from low signal strength. Therefore, it is advantageous for *ex vivo* array coils to provide a high encoding capability for SMS in order to accommodate modern acquisition techniques. Commonly used *in vivo* head coils do not optimally fulfill this requirement for SMS *ex vivo* scans, as they lack enough elements in the *z*-direction. The radially surrounding, *z*-directional, stacked elements of the constructed coil provide favorable spatial coverage for SMS image encoding, allowing the separation of multiple collapsed slices. In the case of an *MB* = 8 acceleration scheme, the combination of the enhanced SMS encoding power and the increased baseline SNR of the 48ch coil, provides an up to 4.5-fold SNR improvement when compared to the 64ch head coil.

Previous work comparing *ex vivo* dMRI to optical imaging suggests that high spatial resolution (1 mm or higher) improves the accuracy of dMRI-derived axonal orientation estimates, and may have a greater impact than high angular resolution or ultra-high *b*-values [15]. That work utilized small human samples that were scanned in a small-bore MRI scanner. The coil presented here paves the way for sub-mm resolution *ex vivo* dMRI on whole human brains at the high *b*-values accessible on the 3T Connectome scanner. This capability will allow us to map the connectional anatomy and microstructure of the human brain at unprecedented resolutions, as well as provide reference data for evaluating *in vivo* dMRI scans to gain deeper insight into human brain structure at multiple scales. We expect this novel coil design, in combination with the current 3T Connectome scanner equipped with 300 mT m^−1^ gradient strengths and next-generation gradient system planned for the Connectome 2.0 project [52], to advance our understanding of human brain circuitry in health and disease.

## 5. Conclusion

A 48ch close-fitting receive array coil for dMRI of whole *ex vivo* human brains at 3T was designed, constructed, and tested with a brain-shaped phantom and an *ex vivo* brain. We characterized the coil with unloaded-to-loaded *Q*-ratio, noise correlation, SNR, SMS G-factor and stability measurements in comparison to a 64ch whole-head *in vivo* coil. Compared to *in vivo* array coils, smaller loop sizes can be used for *ex vivo* brain samples due to increased loading characteristics of the fixed brain tissue. This allows the design of high-channel count arrays, improving both peripheral SNR and encoding performance for accelerated imaging. Due to the high SNR and parallelism, the designed coil is well-suited for high-resolution, high *b*-value *ex vivo* dMRI acquisitions.

## Abbrevations

48ch: 48-channel
64ch: 64-channel
BW: bandwidth
CAD: computer aided design
dMRI: Diffusion MRI
DTI: Diffusion Tensor Imaging
EPI: echo planar imaging
EPROM: Erasable Programmable Read-Only Memory
F: flip angle
FA: fractional anisotropy
FOV: field of view
M: matrix
PC: polycarbonate
PCB: printed circuit board
PD: proton density
PLP: periodate-lysine-paraformaldehyde
RF: radio frequency
ROI: region of interest
SMS: simultaneous multislice
SNR: signal-to-noise-ratio
TE: echo time
TR: repetition time
VNA: vector network analyzer

## Footnotes

1 under development and not commercially available in the U.S. and its future availability cannot be assured.

## Notes

### Competing Interest Statement

The authors have declared no competing interest.

## References

[1] M. Lagana, M. Rovaris, A. Ceccarelli, C. Venturelli, S. Marini, G. Baselli, Dti parameter optimisation for acquisition at 1.5 t: Snr analysis and clinical application, Computational intelligence and neuroscience 2010 (2010) 8. doi:10.1155/2010/254032.

[2] S. Mori, J. Zhang, Principles of diffusion tensor imaging and its applications to basic neuro-science research, Neuron 51 (5) (2006) 527–539. doi:10.1016/j.neuron.2006.08.012.

[3] J. A. McNab, S. Jbabdi, S. C. Deoni, G. Douaud, T. E. Behrens, K. L. Miller, High resolution diffusion-weighted imaging in fixed human brain using diffusion-weighted steady state free precession, Neuroimage 46 (3) (2009) 775–785. doi:10.1016/j.neuroimage.2009.01.008.

[4] H. Okano, P. Mitra, Brain-mapping projects using the common marmoset, Neuroscience re-search 93 (2015) 3–7. doi:10.1016/j.neures.2014.08.014.

[5] T. E. Conturo, N. F. Lori, T. S. Cull, E. Akbudak, A. Z. Snyder, J. S. Shimony, R. C. McKinstry, H. Burton, M. E. Raichle, Tracking neuronal fiber pathways in the living human brain, Proceedings of the National Academy of Sciences 96 (18) (1999) 10422–10427. doi: 10.1073/pnas.96.18.10422.

[6] P. J. Basser, J. Mattiello, D. LeBihan, Mr diffusion tensor spectroscopy and imaging, Biophys-ical journal 66 (1) (1994) 259–267. doi:10.1016/S0006-3495(94)80775-1.

[7] H. Zeng, Mesoscale connectomics, Current opinion in neurobiology 50 (2018) 154–162. doi: 10.1016/j.conb.2018.03.003.

[8] A. Roebroeck, K. L. Miller, M. Aggarwal, Ex vivo diffusion mri of the human brain: Technical challenges and recent advances, NMR in Biomedicine 32 (2019) e3941. doi:https://doi.org/10.1002/nbm.3941.

[9] J. Beaujoin, N. Palomero-Gallagher, F. Boumezbeur, M. Axer, J. Bernard, F. Poupon, D. Schmitz, J.-F. Mangin, C. Poupon, Post-mortem inference of the human hippocampal connectivity and microstructure using ultra-high field diffusion mri at 11.7 t, Brain Structure and Function 223 (5) (2018) 2157–2179. doi:10.1007/s00429-018-1617-1.

[10] M. Modo, T. K. Hitchens, J. R. Liu, R. M. Richardson, Detection of aberrant hippocampal mossy fiber connections: ex vivo mesoscale diffusion mri and microtractography with histological validation in a patient with uncontrolled temporal lobe epilepsy, Human brain mapping 37 (2) (2016) 780–795. doi:10.1002/hbm.23066.

[11] K. L. Miller, C. J. Stagg, G. Douaud, S. Jbabdi, S. M. Smith, T. E. Behrens, M. Jenkinson, S. A. Chance, M. M. Esiri, N. L. Voets, et al., Diffusion imaging of whole, post-mortem human brains on a clinical mri scanner, Neuroimage 57 (1) (2011) 167–181. doi:10.1016/j.neuroimage.2011.03.070.

[12] J. C. Augustinack, K. Helmer, K. E. Huber, S. Kakunoori, L. Zöllei, B. Fischl, Direct visu-alization of the perforant pathway in the human brain with ex vivo diffusion tensor imaging, Frontiers in human neuroscience 4 (2010) 42. doi:10.3389/fnhum.2010.00042.

[13] F. Fritz, S. Sengupta, R. Harms, D. Tse, B. Poser, A. Roebroeck, Ultra-high resolution and multi-shell diffusion mri of intact ex vivo human brains using kt-dsteam at 9.4 t, Neuroimage 202 (2019) 116087. doi:10.1016/j.neuroimage.2019.116087.

[14] J. Mollink, M. Kleinnijenhuis, A.-M. van Cappellen, S. N. Sotiropoulos, M. Cot-taar, C. Mirfin, M. P. Heinrich, M. Jenkinson, M. Pallebage-Gamarallage, O. Ansorge, S. Jbabdi, K. L. Miller, Evaluating fibre orientation dispersion in white matter: Compari-son of diffusion mri, histology and polarized light imaging, NeuroImage 157 (2017) 561–574. doi:https://doi.org/10.1016/j.neuroimage.2017.06.001. URL http://www.sciencedirect.com/science/article/pii/S1053811917304706

[15] R. Jones, G. Grisot, J. Augustinack, C. Magnain, D. A. Boas, B. Fischl, H. Wang, A. Yendiki, Insight into the fundamental trade-offs of diffusion mri from polarization-sensitive optical co-herence tomography in ex vivo human brain, NeuroImage 214 (2020) 116704. doi:https://doi.org/10.1016/j.neuroimage.2020.116704. URL http://www.sciencedirect.com/science/article/pii/S1053811920301919

[16] M. Pallebage-Gamarallage, S. Foxley, R. A. Menke, I. N. Huszar, M. Jenkinson, B. C. Tendler, C. Wang, S. Jbabdi, M. R. Turner, K. L. Miller, et al., Dissecting the pathobiology of altered mri signal in amyotrophic lateral sclerosis: A post mortem whole brain sampling strategy for the integration of ultra-high-field mri and quantitative neuropathology, BMC neuroscience 19 (1) (2018) 11. doi:10.1186/s12868-018-0416-1.

[17] B. R. Plantinga, A. Roebroeck, V. G. Kemper, K. Uludağ, M. Melse, J. Mai, M. L. Kuijf, A. Herrler, A. Jahanshahi, B. M. ter Haar Romeny, et al., Ultra-high field mri post mortem structural connectivity of the human subthalamic nucleus, substantia nigra, and globus pal-lidus, Frontiers in neuroanatomy 10 (2016) 66. doi:10.3389/fnana.2016.00066.

[18] J. C. Augustinack, K. Helmer, K. E. Huber, S. Kakunoori, L. Zöllei, B. Fischl, Direct visu-alization of the perforant pathway in the human brain with ex vivo diffusion tensor imaging, Frontiers in human neuroscience 4 (2010) 42. doi:10.3389/fnhum.2010.00042.

[19] F. Calamante, J.-D. Tournier, N. D. Kurniawan, Z. Yang, E. Gyengesi, G. J. Galloway, D. C. Reutens, A. Connelly, Super-resolution track-density imaging studies of mouse brain: comparison to histology, Neuroimage 59 (1) (2012) 286–296. doi:10.1016/j.neuroimage.2011.07.014.

[20] J. A. McNab, B. L. Edlow, T. Witzel, S. Y. Huang, H. Bhat, K. Heberlein, T. Feiweier, K. Liu, B. Keil, J. Cohen-Adad, et al., The human connectome project and beyond: initial applications of 300 mt/m gradients, Neuroimage 80 (2013) 234–245. doi:10.1016/j.neuroimage.2013.05.074.

[21] K. Setsompop, R. Kimmlingen, E. Eberlein, T. Witzel, J. Cohen-Adad, J. A. McNab, B. Keil, M. D. Tisdall, P. Hoecht, P. Dietz, et al., Pushing the limits of in vivo diffusion mri for the human connectome project, Neuroimage 80 (2013) 220–233. doi:10.1016/j.neuroimage.2013.05.078.

[22] C. Eichner, M. Paquette, T. Mildner, T. Schlumm, K. Pléh, L. Samuni, C. Crockford, R. M. Wittig, C. Jäger, H. E. Möller, A. D. Friederici, A. Anwander, Increased sensitivity and signal-to-noise ratio in diffusion-weighted mri using multi-echo acquisitions, NeuroImage 221 (2020) 117172. doi:https://doi.org/10.1016/j.neuroimage.2020.117172. URL http://www.sciencedirect.com/science/article/pii/S1053811920306583

[23] A. Roebroeck, S. Sengupta, M. Bastiani, S. Schillak, B. Tramm, M. Waks, A. Lataster, A. Her-rler, D. Tse, B. Poser, High resolution mri neuroanatomy of the whole human brain post mortem with a specialized 9.4 t rf-coil, Proc. OHBMdoi:10.13140/RG.2.2.21380.07041.

[24] S. Sengupta, F. Fritz, R. Harms, S. Hildebrand, D. Tse, B. A. Poser, R. Goebel, A. Roebroeck, High resolution anatomical and quantitative mri of the entire human occipital lobe ex vivo at 9.4 t, Neuroimage 168 (2018) 162–171. doi:10.1016/j.neuroimage.2017.03.039.

[25] B. L. Edlow, A. Mareyam, A. Horn, J. R. Polimeni, T. Witzel, M. D. Tisdall, J. C. Augustinack, J. P. Stockmann, B. R. Diamond, A. Stevens, et al., 7 tesla mri of the ex vivo human brain at 100 micron resolution, Scientific data 6 (1) (2019) 1–10. doi:10.1038/s41597-019-0254-8.

[26] A. Scholz, M. May, R. Etzel, M. Mahmutovic, N. Kutscha, L. L. Wald, A. Yendiki, B. Keil, A 48-channel ex vivo brain array coil for diffusion-weighted mri at 3t, in: Proceedings of the 27th Annual Meeting of ISMRM, Montréal, 2019, p. 1494.

[27] G. C. Wiggins, C. Triantafyllou, A. Potthast, A. Reykowski, M. Nittka, L. Wald, 32-channel 3Tesla receive-only phased-array head coil with soccer-ball element geometry, Magnetic Resonance in Medicine: An Official Journal of the International Society for Magnetic Resonance in Medicine 56 (1) (2006) 216–223. doi:10.1002/mrm.20925.

[28] P. B. Roemer, W. A. Edelstein, C. E. Hayes, S. P. Souza, O. M. Mueller, The nmr phased array, Magnetic resonance in medicine 16 (2) (1990) 192–225. doi:10.1002/mrm.1910160203.

[29] A. Kumar, W. A. Edelstein, P. A. Bottomley, Noise figure limits for circular loop mr coils, Magnetic Resonance in Medicine 61 (5) (2009) 1201–1209. doi:10.1002/mrm.21948. URL https://onlinelibrary.wiley.com/doi/abs/10.1002/mrm.21948

[30] B. Keil, L. L. Wald, Massively parallel mri detector arrays, Journal of magnetic resonance 229 (2013) 75–89. doi:10.1016/j.jmr.2013.02.001.

[31] A. Reykowski, S. M. Wright, J. R. Porter, Design of matching networks for low noise preamplifiers, Magnetic resonance in medicine 33 (6) (1995) 848–852. doi:10.1002/mrm.1910330617.

[32] W. Edelstein, C. Hardy, O. Mueller, Electronic decoupling of surface-coil receivers for nmr imaging and spectroscopy, Journal of Magnetic Resonance (1969) 67 (1) (1986) 156–161. doi: 10.1016/0022-2364(86)90421-X.

[33] D. M. Peterson, B. L. Beck, G. R. Duensing, J. R. Fitzsimmons, Common mode signal rejection methods for mri: reduction of cable shield currents for high static magnetic field systems, Concepts in Magnetic Resonance Part B: Magnetic Resonance Engineering: An Educational Journal 19 (1) (2003) 1–8. doi:10.1002/cmr.b.10090.

[34] D. Hoult, The nmr receiver: a description and analysis of design, Progress in Nuclear Magnetic Resonance Spectroscopy 12 (1) (1978) 41–77. doi:10.1016/0079-6565(78)80002-8.

[35] C. Ianniello, J. A. de Zwart, Q. Duan, C. M. Deniz, L. Alon, J.-S. Lee, R. Lattanzi, R. Brown, Synthesized tissue-equivalent dielectric phantoms using salt and polyvinylpyrrolidone solutions, Magnetic resonance in medicine 80 (1) (2018) 413–419. doi:10.1002/mrm.27005.

[36] P. Kellman, E. R. McVeigh, Image reconstruction in snr units: a general method for snr measurement, Magnetic resonance in medicine 54 (6) (2005) 1439–1447. doi:10.1002/mrm.20713.

[37] K. P. Pruessmann, M. Weiger, M. B. Scheidegger, P. Boesiger, Sense: sensitivity encoding for fast mri, Magnetic resonance in medicine 42 (5) (1999) 952–962. doi:10.1002/(SICI)1522-2594(199911)42:5<952::AID-MRM16>3.0.CO;2-S.

[38] D. J. Larkman, J. V. Hajnal, A. H. Herlihy, G. A. Coutts, I. R. Young, G. Ehnholm, Use of multicoil arrays for separation of signal from multiple slices simultaneously excited, Journal of Magnetic Resonance Imaging: An Official Journal of the International Society for Magnetic Resonance in Medicine 13 (2) (2001) 313–317. doi:10.1002/1522-2586(200102)13:2<313::AID-JMRI1045>3.0.CO;2-W.

[39] K. Setsompop, B. A. Gagoski, J. R. Polimeni, T. Witzel, V. J. Wedeen, L. L. Wald, Blipped-controlled aliasing in parallel imaging for simultaneous multislice echo planar imaging with reduced g-factor penalty, Magnetic resonance in medicine 67 (5) (2012) 1210–1224. doi: 10.1002/mrm.23097.

[40] D. A. Feinberg, S. Moeller, S. M. Smith, E. Auerbach, S. Ramanna, M. F. Glasser, K. L. Miller, K. Ugurbil, E. Yacoub, Multiplexed echo planar imaging for sub-second whole brain fmri and fast diffusion imaging, PloS one 5 (12). doi:10.1371/journal.pone.0015710.

[41] B. Keil, J. N. Blau, S. Biber, P. Hoecht, V. Tountcheva, K. Setsompop, C. Triantafyllou, L. L. Wald, A 64-channel 3t array coil for accelerated brain mri, Magnetic resonance in medicine 70 (1) (2013) 248–258. doi:10.1002/mrm.24427.

[42] R. M. Weisskoff, Simple measurement of scanner stability for functional nmr imaging of activation in the brain, Magnetic resonance in medicine 36 (4) (1996) 643–645. doi:10.1002/mrm.1910360422.

[43] J. L. Andersson, S. N. Sotiropoulos, An integrated approach to correction for off-resonance effects and subject movement in diffusion mr imaging, NeuroImage 125 (2016) 1063–1078. doi:https://doi.org/10.1016/j.neuroimage.2015.10.019. URL http://www.sciencedirect.com/science/article/pii/S1053811915009209

[44] B. Keil, V. Alagappan, A. Mareyam, J. A. McNab, K. Fujimoto, V. Tountcheva, C. Triantafyllou, D. D. Dilks, N. Kanwisher, W. Lin, et al., Size-optimized 32-channel brain ar-rays for 3T pediatric imaging, Magnetic Resonance in Medicine 66 (6) (2011) 1777–1787. doi:10.1002/mrm.22961.

[45] T. Janssens, B. Keil, R. Farivar, J. A. McNab, J. R. Polimeni, A. Gerits, J. Arsenault, L. L. Wald, W. Vanduffel, An implanted 8-channel array coil for high-resolution macaque mri at 3T, Neuroimage 62 (3) (2012) 1529–1536. doi:10.1016/j.neuroimage.2012.05.028.

[46] F. Wiesinger, N. De Zanche, K. Pruessmann, Approaching ultimate snr with finite coil arrays, in: Proceedings of the 13th Annual Meeting of ISMRM, 2005, p. 672.

[47] A. S. Shatil, M. N. Uddin, K. M. Matsuda, C. R. Figley, Quantitative ex vivo mri changes due to progressive formalin fixation in whole human brain specimens: longitudinal characterization of diffusion, relaxometry, and myelin water fraction measurements at 3t, Frontiers in medicine 5 (2018) 31. doi:10.3389/fmed.2018.00031.

[48] J. E. Iglesias, R. Insausti, G. Lerma-Usabiaga, M. Bocchetta, K. Van Leemput, D. N. Greve, A. Van der Kouwe, B. Fischl, C. Caballero-Gaudes, P. M. Paz-Alonso, et al., A probabilistic atlas of the human thalamic nuclei combining ex vivo mri and histology, Neuroimage 183 (2018) 314–326. doi:10.1016/j.neuroimage.2018.08.012.

[49] A. S. Shatil, K. M. Matsuda, C. R. Figley, A method for whole brain ex vivo magnetic resonance imaging with minimal susceptibility artifacts, Frontiers in neurology 7 (2016) 208. doi:10.3389/fneur.2016.00208.

[50] S. Y. Huang, Q. Tian, Q. Fan, T. Witzel, B. Wichtmann, J. A. McNab, J. D. Bireley, N. Machado, E. C. Klawiter, C. Mekkaoui, L. L. Wald, A. Nummenmaa, High-gradient diffusion mri reveals distinct estimates of axon diameter index within different white matter tracts in the in vivo human brain, Brain Structure and Function 225 (2020) 1277–1291. doi:10.1007/s00429-019-01961-2.

[51] K. Setsompop, Q. Fan, J. Stockmann, B. Bilgic, S. Huang, S. Cauley, A. Nummenmaa, F. Wang, Y. Rathi, T. Witzel, W. Ll, High-resolution in vivo diffusion imaging of the human brain with generalized slice dithered enhanced resolution: Simultaneous multislice (gslider-sms), Magnetic Resonance in Medicine (2018) 141–151doi:10.1002/mrm.26653.

[52] A. Yendiki, T. Witzel, S. Y. Huang, Connectome 2.0: Cutting-edge hardware ushers in new opportunities for computational diffusion mri, in: E. Bonet-Carne, J. Hutter, M. Palombo, M. Pizzolato, F. Sepehrband, F. Zhang (Eds.), Computational Diffusion MRI, Springer, Switzerland, 2020, pp. 3–12. doi:10.1007/978-3-030-52893-5_1.

